# Clinicopathologic Correlates and Natural History of Atypical Chronic Myeloid Leukemia

**DOI:** 10.1101/2020.09.10.291641

**Authors:** Guillermo Montalban-Bravo, Rashmi Kanagal-Shamanna, Koji Sasaki, Lucia Masarova, Kiran Naqvi, Elias Jabbour, Courtney DiNardo, Koichi Takahashi, Marina Konopleva, Naveen Pemmaraju, Tapan Kadia, Farhad Ravandi, Naval Daver, Gautam Borthakur, Zeev Estrov, Joseph D. Khoury, Sanam Loghavi, Kelly A. Soltysiak, Sherry Pierce, Carlos Bueso-Ramos, Keyur Patel, Srdan Verstovsek, Hagop Kantarjian, Prithviraj Bose, Guillermo Garcia-Manero

**Author notes:** These authors contributed equally to this work (Guillermo Montalban-Bravo, Rashmi Kanagal-Shamanna). **Corresponding Author:** Guillermo Montalban-Bravo, MD, Assistant Professor, The University of Texas MD Anderson Cancer Center, Department of Leukemia, 1515 Holcombe Blvd, Unit 0428, Houston, TX 77030.

## Abstract

There is limited data on the clonal mechanisms underlying leukemogenesis, prognostic factors, and optimal therapy for atypical chronic myeloid leukemia (aCML). We evaluated the clinicopathological features, outcomes, and responses to therapy of 65 patients with aCML. Median age was 67 years (range 46-89). The most frequently mutated genes included *ASXL1* (83%), *SRSF2* (68%), and *SETBP1* (58%). Mutations in *SETBP1, SRSF2, TET2*, and *GATA2* tended to appear within dominant clones, with frequent *SRSF2* and *SETBP1* codominance, while other RAS pathway mutations were more likely to appear as minor clones. Acquisition of new, previously undetectable mutations at transformation was observed in 63% of evaluable patients, the most common involving signaling pathway mutations. Hypomethylating agents were associated with the highest response rates and duration. With a median overall survival of 25 months (95% CI 20-30), intensive chemotherapy was associated with worse OS than other treatment modalities, and allogeneic stem cell transplantation was the only therapy associated with improved outcomes (HR 0.044, 95% CI 0.035-0.593, p=0.007). Age, platelet count, BM blast percentage, and serum LDH levels were independent predictors of survival and were integrated in a multivariable model which allowed to predict 1-year and 3-year survival.

## INTRODUCTION

Atypical chronic myeloid leukemia (aCML) is a rare clonal, hematopoietic stem-cell disorder classified among the myelodysplastic/myeloproliferative neoplasms (MDS/MPN). Atypical CML is characterized by the presence of hypercellular bone marrow (BM) with granulocytic proliferation and granulocytic dysplasia along with peripheral blood (PB) leukocytosis with increased numbers of neutrophils and immature granulocytic precursors comprising ≥10% of leukocytes, in the absence of absolute basophilia and monocytosis and *BCR-ABL1* rearrangement or other features of MPN (1). Next-generation sequencing (NGS) identified recurrent mutations in *ASXL1, SETBP1, ETNK1, TET2*, and other RAS pathway mutations, as well as *CSF3R* mutations, in aCML (2–10). In addition, recent data suggests that co-mutation patterns may be associated with distinct MDS/MPN subtypes, with *ASXL1* and *SETBP1* co-mutations frequently observed in aCML (7). However, the clonal dominance of identified mutations in aCML remains poorly understood and the clonal mechanisms associated with disease progression and transformation have not been well characterized.

Furthermore, aCML is characterized by a short median overall survival of 25 months and high risk of transformation to acute myeloid leukemia (AML) compared to other MDS/MPNs (3, 11), however there is scarce data evaluating the potential factors associated with aCML prognosis (3). To date, therapeutic options for patients with aCML are limited and, although therapy with ruxolitnib can be associated with responses in patients with *CSF3R* mutations (5, 12, 13), particularly in the absence of *SETBP1* mutations, there is insufficient evidence on the optimal therapeutic strategies for these patients. Although several reports including small patient cohorts have described the potential use of hypomethylating agents, such as decitabine or azacitidine, for the treatment of aCML (3, 14–18), evaluation of the survival benefit or clinical activity of these compounds a lager cohort and comparison to other therapeutic approaches is needed.

In order to study the clonal architecture and clinical outcomes of patients with aCML based on therapeutics modality and factors associated with transformation and predictors of outcome, we evaluated a cohort of 65 patients with aCML treated at a single institution.

## MATERIALS AND METHODS

### Patients and Samples

We evaluated all consecutive patients with atypical chronic myeloid leukemia treated at The University of Texas MD Anderson Cancer Center (MDACC) from 2005 to 2020. Informed consent was obtained according to protocols approved by the MDACC institutional review board in accordance with the Declaration of Helsinki. Diagnosis of aCML was confirmed in a hematopathology laboratory at MDACC using the 2016 WHO criteria, by two independent hematopathologists (CBR and RKS) (19). Conventional karyotyping was performed on fresh BM aspirates using standard procedures and reported following ISCN 2013 Nomenclature (20).

### Targeted gene sequencing analysis

Genomic DNA was extracted from whole bone marrow aspirate samples and was subject to targeted PCR-based sequencing using a NGS platform evaluating a total of 81 or 28 genes (21). This analysis was performed within the MDACC CLIA-certified Molecular Diagnostics Laboratory (additional details in Supplemental Methods). For NGS-based analysis, the limit of detection for variant calling was 2%. Previously described somatic mutations registered at the Catalogue of Somatic Mutations in Cancer (COSMIC: http://cancer.sanger.ac.uk/cosmic) were considered as potential driver mutations. Variant allele frequency (VAF) estimates of identified mutations were used to evaluate clonal relationships within each individual patient, with clones with the highest VAF or with VAF close to 0.4 being defined as dominant and those present at VAF <0.2 in the presence of another dominant clone being defined as minor. In addition, PCR-based DNA analysis was performed to detect internal tandem duplications and codon 835/836 point mutations in *FLT3*. Multiplex PCR using fluorescently labeled primers was performed, followed by detection and sizing of PCR products using capillary electrophoresis. For detecting point mutations in codons 835/836, restriction enzyme digestion of the PCR products was performed prior to capillary electrophoresis. The lower limit of detection (analytical sensitivity) of this assay was 1% of mutant DNA in a background of wild-type DNA. The ratio of the area under the peak of mutant over total (mutant: wild-type) *FLT3* was used to determine the mutant allele burden.

### Statistical analysis and response assessment

VAF estimates were used to evaluate clonal relationships within each individual sample (22). Clonal relationships were tested using Pearson goodness-of-fit tests with heterogeneity being defined in cases with goodness-of-fit p-values <0.05. Response outcomes were evaluated following the MDS/MPN IWG response criteria (23) for therapy at the time of aCML diagnosis and following ELN 2017 criteria for therapies at the time of transformation to AML (24). Generalized linear models were used to study the association of overall response (ORR), complete remission (CR), and risk factors. Overall survival (OS) was calculated as the time from diagnosis to death or last follow-up date. Event-free survival (EFS) was calculated from the time of initial therapy until relapse, absence of response, or death. Leukemia-free survival (LFS) was calculated from the time of diagnosis to transformation, death, or last follow-up date. Patients who were alive at their last follow-up were censored on that date. The Kaplan-Meier product limit method (25) was used to estimate the median OS, EFS, and LFS for each clinical/demographic factor. Univariate and multivariate Cox proportional hazards regression analyses were used to identify any association with each of the variables and survival outcomes.

## RESULTS

### Clinical and Histopathological Characteristics

A total of 65 patients with aCML were evaluated during the reviewed time period. Patient characteristics are summarized in Table 1. Median age was 67 years (range 46-89), and median WBC was 44.5×10^9^/L (range 5.9-474.9×10^9^/L). Median percentages of immature granulocytes in peripheral blood were as follows: 0% (range 0-27%) promyelocytes, 0% myelocytes (range 0-35%), and 16% metamyelocytes (range 0-51%). Median hemoglobin was 10.0g/dL (range 5.7-14.7g/dL) and median platelet count was 93×10^9^/L (range 12-560×10^9^/L). Forty-one (63%) patients had normal karyotype, with the most frequent recurrent cytogenetic abnormalities including trisomy 8 in 5 (8%) patients, i(17q) in 2 (3%) patients, and del(20q) in 2 (3%) patients. A total of 3 (5%) patients had complex karyotype defined by presence of more than 3 abnormalities, but only 1 patient had a monosomal karyotype. Median European Cooperative Group performance status at the time of diagnosis was 1 (range 0-4). 21 (32%) patients required transfusions prior to the time of evaluation. Significant palpable splenomegaly was observed in 26 (40%) patients and a total of 7 (11%) had extramedullary disease, either confirmed by histopathological evaluation or highly suspected due to imaging including: pathology proven leukemia cutis in 3, gingival hyperplasia in 1 patient, and lymphadenopathy in 3 patients. Nineteen (29%) patients had required hydroxyurea for control of leukocytosis prior to their presentation at MDACC, with 6 (%) patients presenting with signs of spontaneous tumor lysis syndrome or acute renal dysfunction. Of these, 3 required rapid cytoreduction with cytarabine and 1 required leukophoresis. A total of 51 (80%) received therapy at MDACC, including single agent hypomethylating agent in 19 (29%), hydroxyurea in 8 (12%), a hypomethylating agent in combination with ruxolitinib in 7 (11%), single agent ruxolitinib in 5 (8%), hypomethylating agents in combination with other investigational agents in 5 (8%), induction chemotherapy in 3 (5%), and other investigational agents in 1 (2%) patient. Two patients remained on observation, 1 patient received allogeneic stem cell transplant directly, and 14 (22%) patients continued care outside of MDACC.

**Table 1.** Patient Characteristics.

| Characteristic | aCML (N=65) |
| --- | --- |
|  | N (%) / Median [range] |
| Age (years) | 68 [46-89] |
| Male | 45 (69) |
| WBC ($\times 10^9/L$) | 44.5 [5.9-474.9] |
| Neutrophils (%) | 64 [0-93] |
| Promyelocytes (%) | 0 [0-27] |
| Myelocytes (%) | 0 [0-66] |
| Metamyelocytes (%) | 16 [0-51] |
| Lymphocytes (%) | 8 [0-63] |
| Monocytes (%) | 2 [0-13] |
| Basophil (%) | 0 [0-6] |
| Eosinophil (%) | 1 [0-15] |
| Blast (%) | 1 [0-16] |
| Hgb (g/dL) | 10 [5.7-14.7] |
| Platelets ($\times 10^9/L$) | 93 [12-560] |
| BM blast (%) | 2 [0-16] |
| BM progranulocytes (%) | 1 [0-12] |
| BM myelocytes (%) | 16 [2-41] |
| BM metamyelocytes (%) | 5 [5-31] |
| BM granulocytes (%) | 36 [10-77] |
| BM basophils (%) | 0 [0-20] |
| BM eosinophils (%) | 1 [0-11] |
| BM monocytes (%) | 2 [0-10] |
| Cytogenetics |  |
| Normal | 41 (63) |
| Trisomy 8 | 5 (8) |
| i(17q) | 2 (3) |
| Monosomy Y | 1 (2) |
| Del(20q) | 2 (3) |
| Complex | 3 (5) |
| Other | 9 (14) |
| NA | 2 (3) |
| Therapy related (%) | 1 (2) |
| ECOG Performance status |  |
| 0-1 | 45 (69) |
| ≥2 | 12 (18) |
| Prior transfusions | 21 (32) |
| Splenomegaly | 26 (40) |
| Extramedullary disease | 7 (11) |
| B symptoms | 16 (25) |

Bone marrow evaluation revealed a markedly hypercellular marrow in all patients with granulocytic proliferation and granulocytic dysplasia. Significant dyserythropoiesis was observed in 26 (40%) patients with dysmegakaryopoiesis being observed in 43 (66%) patients. Marked trilineage dysplasia was apparent in 22 (33%) patients. Bone marrow grading of fibrosis was performed on a total of 52 (80%) patients with 7 (13%) patients having MF-0, 32 (62%) MF-1, 11 (21%) MF-2, and 2 (4%) MF-3. Median bone myeloid population frequencies are detailed in Table 2.

**Table 2.** Response outcomes in patients with aCML based on therapy.

| **Response** | **HMA**<br>**(n=19)**<br>**N (%)/[range]** | **HMA+ruxolitinib**<br>**(n=6)**<br>**N (%)/[range]** | **Ruxolitinib**<br>**(n=5)**<br>**N**<br>**(%)/[range]** | **HMA-combo**<br>**(n=4)**<br>**N (%)/[range]** | **Chemotherapy**<br>**(n=3)**<br>**N (%)/[range]** |
| --- | --- | --- | --- | --- | --- |
| Overall response rate | 5 (26) | 0 (0) | 1 (9) | 2 (50) | 3 (100) |
| Complete response (CR) | 3 (16) | 0 (0) | 0 (0) | 0 (0) | 0 (0) |
| Marrow CR | 0 (0) | 0 (0) | 0 (0) | 0 (0) | 3 (100) |
| Partial response | 0 (0) | 0 (0) | 0 (0) | 1 (25) | 0 (0) |
| Symptom response | 0 (0) | 0 (0) | 1 (9) | 0 (0) | 0 (0) |
| Clinical benefit | 2 (11) | 0 (0) | 0 (0) | 1 (25) | 0 (0) |
| Median number of cycles | 5 [2-7] | - | - | 5 [2-7] | 2 [1-3] |
| Median number of cycles to best response | 2 [1-5] | - | - | 3 [2-3] | 1 [0-1] |
| Median response duration (months)* | 2.7 [1.9-5.2] | - | *0.4 months | 1.9 [0-3.8] | 2.2 [1-2.5] |
| No response | 10 (53) | 5 (83) | 3 (60) | 2 (50) | 0 (0) |
| Progressive disease | 4 (21) | 1 (17) | 1 (20) | 0 (0) | 0 (0) |

\*Response duration was censored to the date of transplant in patients who underwent allogeneic SCT.
\*: response duration corresponding to a single patient who responded to therapy.

### Mutation and clonal landscape and clinicopathological associations

Next-generation sequencing data was available for 35 (54%) patients. The median number of detectable mutations was 4 (range 1-8). The most frequently mutated genes included *ASXL1* in 83%, *SRSF2* in 68%, and *SETBP1* in 58%. The frequencies of identified mutations are shown in Figure 1A. Mutations in *ASXL1* included frameshift (n=25) or nonsense (n=4) mutations, the most common of which being G646fs in 17/29 patients. The most frequent *SETBP1* mutation included D868N. Other genes with mutations present at a frequency >10% of the evaluated population included *TET2, CBL, GATA2, NRAS, RUNX1, NF1*, and *EZH2*. The median variant allele frequencies of identified mutations are shown in Figure 1B.

**Figure 1.**
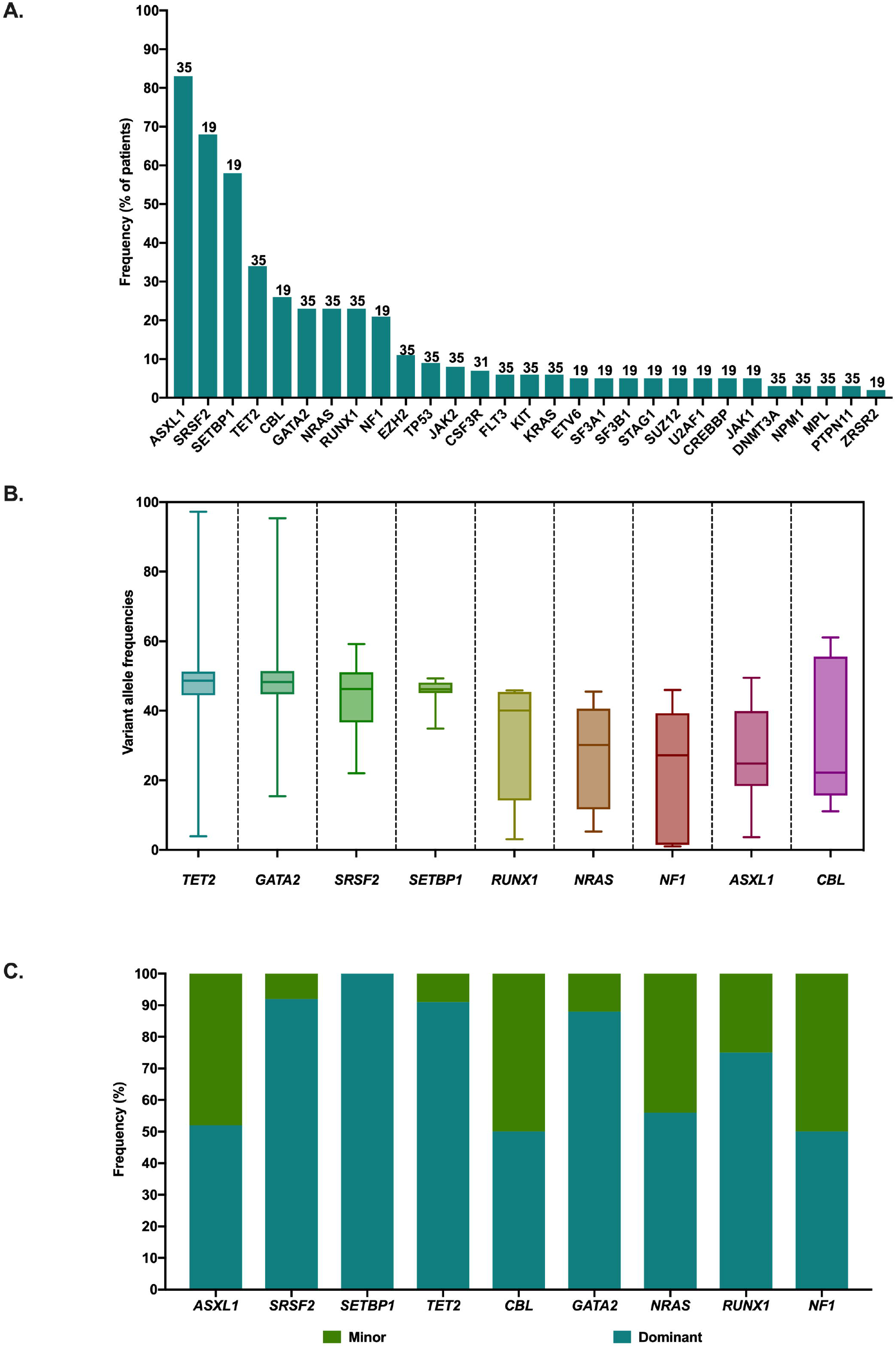
Mutational and clonal landscape of aCML. **A.** Frequency of identified mutations. Number above each specific gene column represents number of patients sequenced for each specific gene. **B.** Median and range of variant allele frequencies (VAFs) of mutations identified in at least 10% of patients. Mutations are ordered by decreasing median VAF. **C.** Frequency of mutations appearing as dominant or minor events. VAF estimates were used to evaluate clonal relationships within each individual sample using Pearson goodness of fit tests and VAF differences. Clones with the highest VAF or with VAFs close to 40% were defined as dominant, and those present at VAF <20% in the presence of another dominant clone were defined as minor.

In order to determine the likely clonal dominance of identified mutations, VAF estimates were used to evaluate clonal relationships within each individual sample (22) using Pearson goodness-of-fit tests and VAF differences. Clones with the highest VAFs or with VAFs close to 40% were defined as dominant, and those present at VAF <20% in the presence of another dominant clone were defined as minor. Mutations in *SETBP1, SRSF2, TET2*, and *GATA2* tended to appear within dominant clones while other RAS pathway mutations were more likely to appear as minor clones. Within the observed commonly co-mutated genes, *SRSF2* and *SETBP1* tended to appear as co-dominant, while *ASXL1*, although the most frequently detected mutation, appeared as a minor clone in up to 50% of patients (Figure 1C).

### Cytogenetic and clonal evolution associated with transformation to acute myeloid leukemia

A total of 18 (28%) patients transformed to AML with a median time to transformation of 18 months (1-123 months). Peripheral blood and bone marrow findings at the time of transformation are detailed in Supplemental Table S1.

Sequencing data at the time of transformation was available for 12 (67%) patients, with matched sequencing at diagnosis of aCML and AML in 8 (44%) patients. The mutational landscape at the time of transformation is shown in Figure 2A. Acquisition of new, previously undetectable mutations was observed in 5 patients, the most common involving signaling pathway mutations (*NRAS, KRAS, NF1, PTPN11*) as well as *FLT3-ITD, ASXL1, CEBPA*, and *ETV6* (Figure 2A). Acquisition of new cytogenetic abnormalities was observed in 9/14 patients, the most frequent involving i(17q). Dynamic changes in the clonal and cytogenetic landscape and disease phenotype during the course of therapy from diagnosis to transformation are shown in Figure 2B.

**Figure 2.**
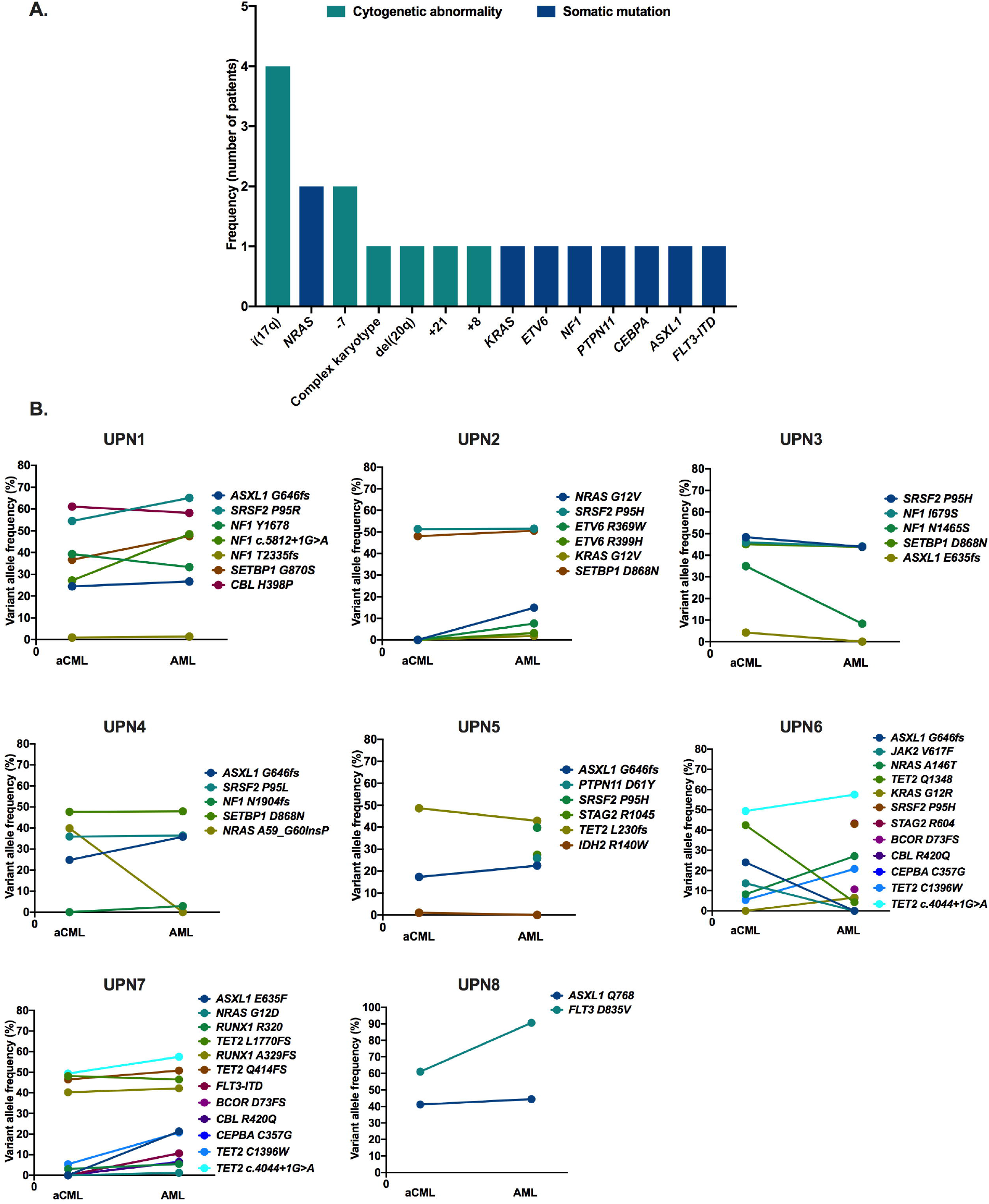
Clonal changes at the time of leukemic transformation. **A.** Frequencies of recurrent somatic mutations and cytogenetic abnormalities identified at the time of leukemic transformation that were not present at diagnosis of aCML. **B.** Dynamic changes in identified somatic mutations and their VAFs from the time of aCML diagnosis to AML.

### Clinical outcomes based on therapy type and genomic and clinical characteristics

With a median follow up of 35.6 months (95% CI 28.2-43.1) a total of 38 (95%) of the patients who received disease-modifying agents were evaluable for response. Patients who continued observation or cytoreductive therapy with hydroxyurea were not considered evaluable for response. Among response-evaluable patients, 19 (50%) received a single agent HMA, 6 (16%) an HMA in combination with ruxolitinib, 5 (13%) single agent ruxolitinib, 4 (11%) an HMA in combination with other investigational agents, 3 (8%) induction chemotherapy, and 1 (3%) proceeded directly to allogeneic stem -cell transplantation. The ORR was 29%, with a total of 3 (8%) patients achieving CR. Response rates and median response durations based on therapeutic modality are detailed in Table 2. Among patients who received ruxolitinib, either as single agent or in combination with an HMA, 2 (17%) had detectable *JAK2* V617F mutations, and no patients had detectable *CSF3R* mutations.

The median OS of the entire cohort was 25 months (95% CI 20.0-30.0, Figure 3A). When evaluating survival based on therapeutic regimen, patients who received intensive chemotherapy had significantly worse OS than those receiving HMA-based therapy or other agents such as ruxolitinib or hydroxyurea (p=0.012, Figure 3B). Of note, among the 3 patients treated with intensive chemotherapy, 1 presented with an ECOG performance status of 4, WBC of 207×10^9^/L, and spontaneous tumor lysis syndrome with acute renal dysfunction at the time of diagnosis, 1 had leukemia cutis with a WBC of 25.8×10^9^/L, and one had gingival hyperplasia and a WBC of 181.4×10^9^/L. No significant differences in survival were observed between patients receiving hydroxyurea, ruxolitinib, or an HMA alone or with other agents. A total of 7 (11%) patients underwent allogeneic stem cell transplantation. The median LFS was 19.8 months (95% CI 15.6-24 months, Figure 3C) and the median survival after transformation was 8.3 months (95% CI 5.5-11.0 months). After transformation to AML 11 patients received therapy with an ORR of 64%, including 4 (36%) CRis and a CR rate of 18%. Therapies included: cladribine or clofarabine in combination with low dose cytarabine (LDAC) in 3 patients; LDAC in combination with venetoclax in 1; an HMA in combination with ruxolitinib in 1; an HMA in combination with venetoclax in 1; an HMA in combination with other agents in 1; intensive chemotherapy with sorafenib in 1; investigational agents in 1; and myeloablative conditioning and transplant in 1. The median number of cycles of therapy was 2 (range 1-5) with a median number of cycles to best response of 2 (range 1-5). Median response duration was 1.4 months (range 0-4) and 2 patients were able to transition to allogeneic stem cell transplant. Among patients who suffered transformation to AML, only 1 remains alive at the time of data cutoff and analysis.

**Figure 3.**
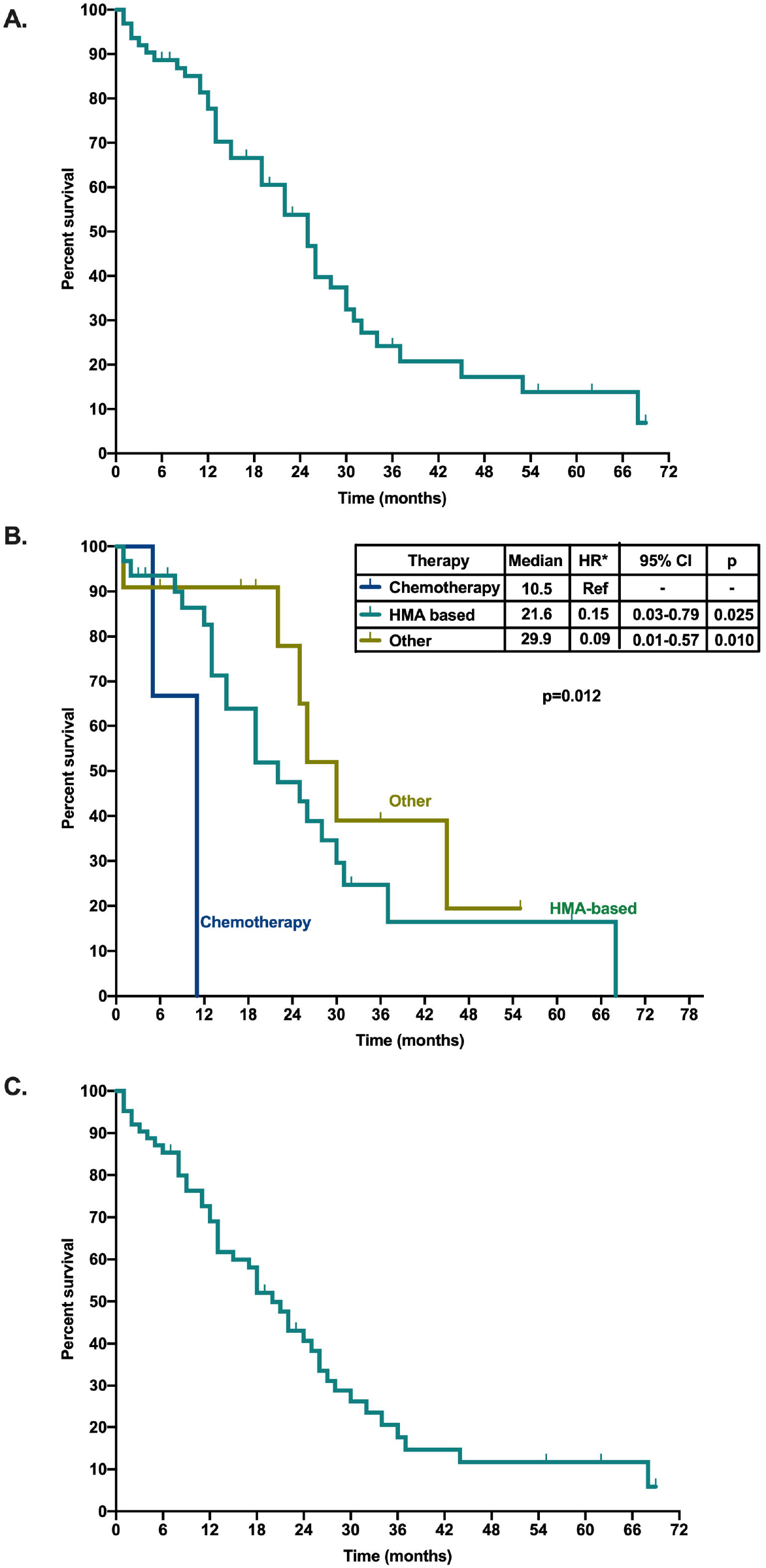
Survival outcomes of patients with aCML. **A.** Kaplan-Meier estimate curve for overall survival of patients with aCML. **B.** Kaplan-Meier estimate curves for overall survival based on type of therapeutic modality. Other includes hydroxyurea or single agent ruxolitinib. **C.** Kaplan-Meier estimate curve for leukemia-free survival of patients with aCML.

By univariate analysis for survival, peripheral blood promyelocyte percentage (p=0.005), performance status ≥2 (p=0.059), hemoglobin (p=0.033), bone marrow blast percentage (p=0.013), and bone marrow monocyte percentage (p=0.027) were associated with survival (Supplemental Table S2). By multivariate analysis for overall survival, age (HR 1.107, 95% CI 1.045-1.173, p=0.001), hemoglobin (HR 0.784, 95% CI 0.635-0.968, p=0.024), platelet count (HR 0.993, 95% CI 0.988-0.997, p=0.003), bone marrow blast percentage (HR 1.414, 95% CI 1.223-1.635, p<0.001), bone marrow monocyte percentage (HR 1.215, 95% CI 1.008-1.466, p=0.041), LDH levels (HR 1.000, 95% CI 1.000-1.000, p<0.001), and allogeneic stem -cell transplantion (HR 0.044, 95% CI 0.035-0.593, p=0.007) were independent predictors of survival (Supplemental Table S2). In order to evaluate disease and patient related features that could allow prediction of the clinical outcomes of patients with aCML at the time of diagnosis, we performed multivariate analysis for survival based on baseline clinicopathological features. The following patient characteristics were independently associated with patient prognosis (Table 3): age, platelet count, bone marrow blast percentage, and LDH levels. This model was used to generate a nomogram for overall survival (Figure 4). This nomogram provides a visual depiction of the relative contribution of each prognostic factor to the total point score and the weight of factors influencing survival. The formula for calculating the total point score is as follows: age points (+60.18185 + 1.337374 x age) + platelet points (59.82308 + −0.99705 x platelet count) + bone marrow blast points (4.54877 x bone marrow blast %) + LDH points (0.002500 x LDH level). Total point scores ranged from 36.1 to 165.1, with a median of 95.0.

**Figure 4.**
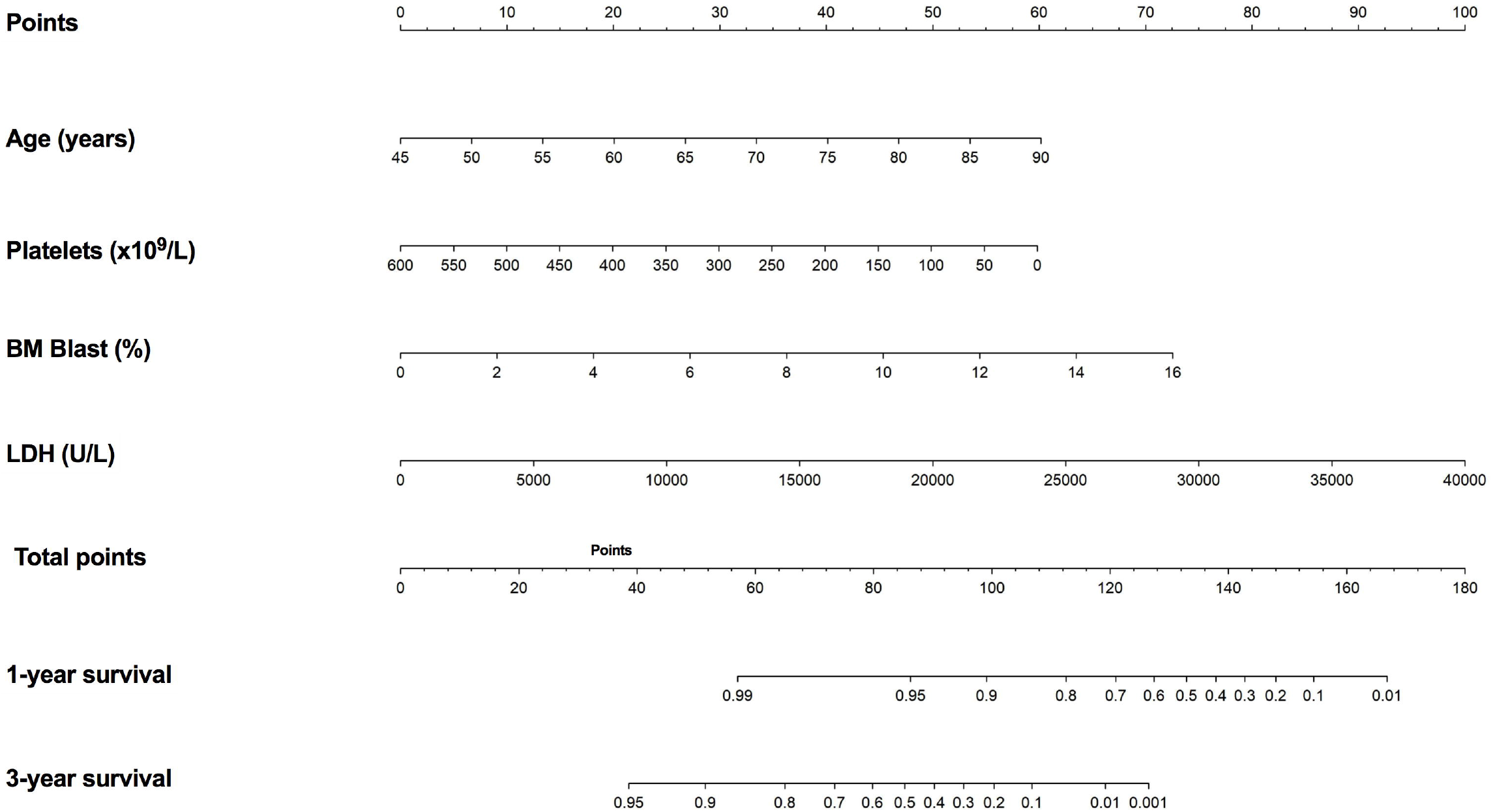
Multivariate Cox proportional hazards model and nomogram for overall survival. Nomogram used by totaling points identified at top scale for each of the independent variables. This summed point score was then identified on a total point scale to identify the 1-year and 3-year survival probabilities.

**Table 3.** Univariate and multivariate analysis for survival based on baseline clinicopathologic features.

|  | UVA |  |  | Backward MVA |  |  |
| --- | --- | --- | --- | --- | --- | --- |
|  | P-value | HR | 95% CI | P value | HR | 95% CI |
| Age | 0.176 | 1.029 | 0.987-1.073 | 0.005 | 1.076 | 1.023-1.133 |
| Female | 0.800 | 0.911 | 0.444-1.872 |  |  |  |
| PS ≥2 | 0.059 | 2.053 | 0.973-4.332 |  |  |  |
| Prior malig. | 0.534 | 1.319 | 0.551-3.153 |  |  |  |
| Prior chemo/XRT | 0.666 | 1.299 | 0.396-4.259 |  |  |  |
| Prior transfusion | 0.551 | 1.233 | 0.620-2.452 |  |  |  |
| Splenomegaly | 0.761 | 1.108 | 0.574-2.139 |  |  |  |
| B symptoms | 0.483 | 1.290 | 0.633-2.630 |  |  |  |
| TLS | 0.876 | 1.087 | 0.383-3.082 |  |  |  |
| WBC | 0.282 | 1.002 | 0.999-1.005 |  |  |  |
| Neu% | 0.498 | 0.994 | 0.977-1.011 |  |  |  |
| Blasts% | 0.192 | 1.051 | 0.975-1.132 |  |  |  |
| Mono% | 0.451 | 0.962 | 0.871-1.063 |  |  |  |
| Lymph% | 0.223 | 0.975 | 0.935-1.016 |  |  |  |
| Baso% | 0.860 | 1.027 | 0.762-1.385 |  |  |  |
| Eo% | 0.076 | 0.849 | 0.708-1.018 |  |  |  |
| Hgb | 0.033 | 0.846 | 0.726-0.987 |  |  |  |
| Plt | 0.156 | 0.998 | 0.995-1.001 | 0.002 | 0.993 | 0.989-0.997 |
| BM blasts | 0.013 | 1.089 | 1.018-1.165 | <0.001 | 1.338 | 1.182-1.515 |
| BM Eo | 0.339 | 0.932 | 0.807-1.077 |  |  |  |
| BM baso | 0.935 | 1.005 | 0.884-1.143 |  |  |  |
| BM mono | 0.027 | 1.172 | 1.018-1.350 |  |  |  |
| EPO | 0.804 | 1.000 | 0.999-1.001 |  |  |  |
| LDH | 0.065 | 1.000 | 1.000-1.000 | <0.001 | 1.000 | 1.000-1.000 |
| UA | 0.435 | 1.042 | 0.939-1.157 |  |  |  |
| Diploid | 0.130 | 0.607 | 0.318-1.158 |  |  |  |

## DISCUSSION

Atypical chronic myeloid leukemia (aCML) is a rare hematopoietic stem cell disorder with dismal prognosis and a high rate of transformation to acute leukemia (11). Although prior reports have described activity of ruxolitinib (12), hydroxyurea, low-dose cytarabine, or HMAs (15, 16, 26) in this disease, data on the optimal clinical management of these patients remains unclear. In addition, given the rarity of this disorder, there is a lack of validated clinical risk models to effectively stratify patients based on predicted outcomes (3, 26). Finally, although several studies have described recurrent somatic mutations in aCML, the clonal architecture in aCML and the genomic changes associated with transformation remain unclear. In this study, we evaluated the clinicopathological features, outcomes, and clonal architecture of a cohort of 65 patients with aCML. By doing so, we observed a high frequency of *SRSF2* and *SETBP1* mutation co-dominance, with *ASXL1* mutations being the most frequently observed; acquisition or clonal expansion of previously undetected RAS pathway mutations; and certain cytogenetic abnormalities, such as i(17q) or monosomy 7, associated with acute transformation. This is consistent with prior reports by our group associating i(17q) with transformation to AML in MDS/MPNs (27). Finally, we developed a prognostic model which included age, platelet count, bone marrow blast percentage, and LDH, which allowed us to predict the survival of patients with aCML.

Prior studies have reported high frequencies of *ASXL1* and *SETBP1* mutations in aCML. In a recently published study by Palomo, *et al* (7), mutations in *ASXL1* strongly correlated with *SETBP1* mutations in patients with aCML. Although in this study the authors identified *ASXL1* mutations as part of ancestral clones in a majority of patients (79%), we identified that both *SRSF2* and *SETBP1* mutations tended to appear at significantly higher VAFs and as dominant events in a majority of patients, while *ASXL1* mutations appeared in minor clones in up to 50% of patients. In addition, similar to their findings, we observed that *GATA2* mutations tended to appear within dominant clones and *RAS* pathway signaling gene mutations tended to appear in minor clones. However, sequential targeted sequencing performed at the time of transformation revealed that, although initially present within non-dominant clones, mutations in signaling genes other than *SETBP1* were associated with clonal expansion and transformation to AML. This data suggests that, although other RAS pathway signaling mutations might be less common and appear as subclonal events in aCML, their expansion through the course of therapy or their acquisition in initial founder clones may likely be responsible for transformation and resistance to therapies. This might be relevant when considering future therapeutic combinations with agents such as BCL2 inhibitors, given the known association of these mutations with resistance to therapies such as venetoclax (28). In addition, this underscores the importance of developing effective agents targeting RAS signaling or MCL-1 in the current era of venetoclax-based therapies (29, 30).

Although prior reports have suggested that therapy with single agent ruxolitinib might be effective in patients with aCML, we did not observe significant responses. However, none of the 5 patients who received single agent ruxolitinib in our cohort had *CSF3R* mutations and only 2 had *JAK2* mutations. In addition, the combination of azacitidine with ruxolitinib was not associated with any significant responses. Prior case reports described the potential efficacy of azacitidine or decitabine in patients with aCML. In our study, response rates to HMA-based therapies were observed in 25% of patients, and only 3 patients achieved CR, with a median response duration of 2.7 months. Although use of intensive chemotherapy was associated with worse survival, patients treated with this therapeutic modality had highly proliferative disease with extramedullary involvement and spontaneous tumor lysis, suggesting that their underlying disease biology was likely responsible for the shorter survival times. Allogeneic stem cell transplantation was the only therapeutic strategy that was associated with significantly improved survival, suggesting that all patients who are eligible should be considered for transplant.

Prior studies evaluating a cohort of 55 and 25 patients with aCML reported advanced age, high WBC, low hemoglobin, presence of immature granulocytic precursors, and *TET2* mutations to be associated with worse survival in patients with aCML (3, 26). In our study, with one of the largest cohorts so far reported, blood immature granulocyte percentage and bone marrow monocytosis were associated with worse survival by univariate analysis, but lost their independent prognostic significance in multivariate analysis, while age, platelet count, bone marrow blast percentage, and LDH levels remained independent predictors of survival. Integration of these variables into a multivariate Cox regression model allowed us to create a nomogram that predicted 1-year and 3-year overall survival. Although further validation of this model is warranted, its integration into clinical practice may allow more specific survival estimates when compared to conventional cutoff driven scoring systems.

We acknowledge that our study has several limitations. First, its retrospective nature limits our ability to unequivocally confirm survival and response differences based on distinct therapeutic interventions. Second, the absence of NGS in all evaluated patients limited our ability to incorporate somatic mutation data as part of the multivariate prognostic model, and only a subset of patients who transformed to AML had sequencing at the time of progression. Finally, although this study includes one of the largest reported clinically annotated cohorts of patients with aCML, prospective studies will be necessary to confirm the optimal therapeutic modality for patients with aCML.

Despite these limitations, our data suggest that aCML is characterized by specific mutational clonal dominance with a high frequency of co-dominant *SRSF2* and *SETBP1* mutations, and other RAS pathway mutations present in minor clones at the time of diagnosis but associated with AML transformation. In addition, we observed that acquisition of i(17q) is associated with AML transformation. Also, we confirm poor survival and response outcomes with most treatment modalities, with HMA treatment associated with the highest and most durable responses. Finally, incorporation of age, platelet count, bone marrow blast percentage, and LDH levels can allow survival prediction for these patients, and allogeneic stem cell transplantation should be considered on all eligible patients with a diagnosis of aCML.

## Data Availability Statement

The datasets generated during and/or analyzed during the current study are not publicly available due to patient privacy concerns but are available from the corresponding author on reasonable request.

## ACKNOWLEDGEMENTS

This work was supported in part by the University of Texas MD Anderson Cancer Center Support Grant CA016672 and the University of Texas MD Anderson Cancer Center MDS/AML Moon Shot.

## REFERENCES

1. Arber DA, Orazi A, Hasserjian R, Thiele J, Borowitz MJ, Le Beau MM, et al. The 2016 revision to the World Health Organization classification of myeloid neoplasms and acute leukemia. Blood. 2016;127(20):2391–405.

2. Makishima H, Yoshida K, Nguyen N, Przychodzen B, Sanada M, Okuno Y, et al. Somatic SETBP1 mutations in myeloid malignancies. Nature genetics. 2013;45(8):942–6.

3. Patnaik MM, Barraco D, Lasho TL, Finke CM, Reichard K, Hoversten KP, et al. Targeted next generation sequencing and identification of risk factors in World Health Organization defined atypical chronic myeloid leukemia. Am J Hematol. 2017;92(6):542–8.

4. Langabeer SE, Comerford CM, Quinn J, Murphy PT. Molecular profiling and targeted inhibitor therapy in atypical chronic myeloid leukaemia in blast crisis. J Clin Pathol. 2017;70(12):1089.

5. Ammatuna E, Eefting M, van Lom K, Kavelaars FG, Valk PJ, Touw IP. Atypical chronic myeloid leukemia with concomitant CSF3R T618I and SETBP1 mutations unresponsive to the JAK inhibitor ruxolitinib. Ann Hematol. 2015;94(5):879–80.

6. Gambacorti-Passerini CB, Donadoni C, Parmiani A, Pirola A, Redaelli S, Signore G, et al. Recurrent ETNK1 mutations in atypical chronic myeloid leukemia. Blood. 2015;125(3):499–503.

7. Palomo L, Meggendorfer M, Hutter S, Twardziok S, Adema V, Fuhrmann I, et al. Molecular landscape and clonal architecture of adult myelodysplastic/myeloproliferative neoplasms. Blood. 2020.

8. Maxson JE, Gotlib J, Pollyea DA, Fleischman AG, Agarwal A, Eide CA, et al. Oncogenic CSF3R mutations in chronic neutrophilic leukemia and atypical CML. N Engl J Med. 2013;368(19):1781–90.

9. Yun JW, Yoon J, Jung CW, Lee KO, Kim JW, Kim SH, et al. Next-generation sequencing reveals unique combination of mutations in cis of CSF3R in atypical chronic myeloid leukemia. J Clin Lab Anal. 2020;34(2):e23064.

10. Zhang H, Wilmot B, Bottomly D, Dao KT, Stevens E, Eide CA, et al. Genomic landscape of neutrophilic leukemias of ambiguous diagnosis. Blood. 2019;134(11):867–79.

11. Wang SA, Hasserjian RP, Fox PS, Rogers HJ, Geyer JT, Chabot-Richards D, et al. Atypical chronic myeloid leukemia is clinically distinct from unclassifiable myelodysplastic/myeloproliferative neoplasms. Blood. 2014;123(17):2645–51.

12. Dao KT, Gotlib J, Deininger MMN, Oh ST, Cortes JE, Collins RH, Jr., et al. Efficacy of Ruxolitinib in Patients With Chronic Neutrophilic Leukemia and Atypical Chronic Myeloid Leukemia. J Clin Oncol. 2020;38(10):1006–18.

13. Dao KH, Solti MB, Maxson JE, Winton EF, Press RD, Druker BJ, et al. Significant clinical response to JAK1/2 inhibition in a patient with CSF3R-T618I-positive atypical chronic myeloid leukemia. Leuk Res Rep. 2014;3(2):67–9.

14. Andriani A, Elli E, Trape G, Villiva N, Fianchi L, Di Veroli A, et al. Treatment of Philadelphia-negative myeloproliferative neoplasms in accelerated/blastic phase with azacytidine. Clinical results and identification of prognostic factors. Hematol Oncol. 2019;37(3):291–5.

15. Andriani A, Montanaro M, Voso MT, Villiva N, Ciccone F, Andrizzi C, et al. Azacytidine for the treatment of retrospective analysis from the Gruppo Laziale for the study of Ph-negative MPN. Leuk Res. 2015;39(8):801–4.

16. Hausmann H, Bhatt VR, Yuan J, Maness LJ, Ganti AK. Activity of single-agent decitabine in atypical chronic myeloid leukemia. J Oncol Pharm Pract. 2016;22(6):790–4.

17. Marumo A, Mizuki T, Tanosaki S. Atypical chronic myeloid leukemia achieving good response with azacitidine. Indian J Cancer. 2019;56(4):354–5.

18. Tong X, Li J, Zhou Z, Zheng D, Liu J, Su C. Efficacy and side-effects of decitabine in treatment of atypical chronic myeloid leukemia. Leukemia & lymphoma. 2015;56(6):1911–3.

19. Arber DA, Orazi A, Hasserjian R, Thiele J, Borowitz MJ, Le Beau MM, et al. The 2016 revision to the World Health Organization (WHO) classification of myeloid neoplasms and acute leukemia. Blood. 2016.

20. Simons A, Shaffer LG, Hastings RJ. Cytogenetic Nomenclature: Changes in the ISCN 2013 Compared to the 2009 Edition. Cytogenet Genome Res. 2013;141(1):1–6.

21. Lindsley RC, Mar BG, Mazzola E, Grauman PV, Shareef S, Allen SL, et al. Acute myeloid leukemia ontogeny is defined by distinct somatic mutations. Blood. 2015;125(9):1367–76.

22. Papaemmanuil E, Gerstung M, Malcovati L, Tauro S, Gundem G, Van Loo P, et al. Clinical and biological implications of driver mutations in myelodysplastic syndromes. Blood. 2013;122(22):3616–27; quiz 99.

23. Savona MR, Malcovati L, Komrokji R, Tiu RV, Mughal TI, Orazi A, et al. An international consortium proposal of uniform response criteria for myelodysplastic/myeloproliferative neoplasms (MDS/MPN) in adults. Blood. 2015;125(12):1857–65.

24. Dohner H, Estey E, Grimwade D, Amadori S, Appelbaum FR, Buchner T, et al. Diagnosis and management of AML in adults: 2017 ELN recommendations from an international expert panel. Blood. 2017;129(4):424–47.

25. Kaplan El MP. Nonparametric estimation from incomplete observations. Journal of the American Statistical Association. 1958(53):457–81.

26. Breccia M, Biondo F, Latagliata R, Carmosino I, Mandelli F, Alimena G. Identification of risk factors in atypical chronic myeloid leukemia. Haematologica. 2006;91(11):1566–8.

27. Kanagal-Shamanna R, Bueso-Ramos CE, Barkoh B, Lu G, Wang S, Garcia-Manero G, et al. Myeloid neoplasms with isolated isochromosome 17q represent a clinicopathologic entity associated with myelodysplastic/myeloproliferative features, a high risk of leukemic transformation, and wild-type TP53. Cancer. 2012;118(11):2879–88.

28. DiNardo CD, Tiong IS, Quaglieri A, MacRaild S, Loghavi S, Brown FC, et al. Molecular patterns of response and treatment failure after frontline venetoclax combinations in older patients with AML. Blood. 2020;135(11):791–803.

29. Hormi M, Birsen R, Belhadj M, Huynh T, Cantero Aguilar L, Grignano E, et al. Pairing MCL-1 inhibition with venetoclax improves therapeutic efficiency of BH3-mimetics in AML. Eur J Haematol. 2020.

30. Tambe M, Karjalainen E, Vaha-Koskela M, Bulanova D, Gjertsen BT, Kontro M, et al. Pan-RAF inhibition induces apoptosis in acute myeloid leukemia cells and synergizes with BCL2 inhibition. Leukemia. 2020.

