## Supplemental Data for "Clinicopathologic Correlates and Natural History of Atypical Chronic Myeloid Leukemia"

**Table of Contents**

**Content	Page**

Supplementary Methods

PCR-based NGS 81-gene panel	2

Supplementary Tables

Table S1. Characteristics at transformation to acute myeloid leukemia	4

Table S2. Univariate and multivariate analysis for overall survival	5

**PCR-based next generation sequencing (NGS) 81-gene panel**

| ***Gene*** | **Exons (codons) tested** |
| --- | --- |
| ***ANKRD26*** | 1 (1-6) |
| ***ASXL1*** | 11-12 (362-1442), 12 (1450-1542) |
| ***ASXL2*** | 11-12 (381-1436) |
| ***BCOR*** | 2-4 (1-511), 4-12 (515-1547), 13-15 (1550-1644), 15 (1663-1722) |
| ***BCORL1*** | 1-6 (1-1261), 6 (1292-1323), 6-12 (1326-1700), 12 (1706-1712) |
| ***BRAF*** | 11 (439-478), 15 (581-620) |
| ***BRINP3*** | 2-8 (1-767) |
| ***CALR*** | 9 (352-418) |
| ***CBL*** | 7-9 (336-477) |
| ***CBLB*** | 7-10 (282-469) |
| ***CBLC*** | 7-9 (336-454), 10 (465-475) |
| ***CEBPA*** | 1 (1-90), 1 (249-358), 1 (128-175), 1 (178-201) |
| ***CREBBP*** | 1-8 (1-608), 9-30 (615-1724), 31 (2238-2443), 31 (2049-2235), 31 (1725-1943), 31 (1950-2042) |
| ***CRLF2*** | 6 (217-256) |
| ***CSF3R*** | 14 (575-622), 17 (681-800), 17 (822-864) |
| ***CUX1*** | 2-6 (11-172), 6-9 (174-241), 10-14 (248-408) |
| ***DDX41*** | 1-11 (1-410), 12 (416-418), 12-17 (420-623) |
| ***DNMT3A*** | 8-22 (286-862), 23 (866-913) |
| ***EED*** | 1-2 (1-69), 2-8 (71-287), 9-12 (289-442) |
| ***ELANE*** | 1-2 (1-48), 2 (69-75), 4-5 (123-268) |
| ***ETNK1*** | 3 (228-275) |
| ***ETV6*** | 1-8 (1-453) |
| ***EZH2*** | 2-5 (1-158), 5-6 (160-205), 7 (209-217), 8-19 (243-732), 20 (752) |
| ***FBXW7*** | 9-12 (413-708) |
| ***FLT3*** | 11-20 (437-847) |
| ***GATA1*** | 2-3 (1-84) |
| ***GATA2*** | 2 (2-5), 2-5 (22-377), 5-6 (379-481) |
| ***GFI1*** | 2 (2-39) |
| ***GNAS*** | 8 (200-202), 11 (315-324) |
| ***HNRNPK*** | 3-17 (1-465) |
| ***HRAS*** | 2-3 (1-59), 3-4 (87-135), 4 (137-150) |
| ***IDH1*** | 4 (132-133) |
| ***IDH2*** | 4 (125-178) |
| ***IKZF1*** | 2-8 (1-443), 8 (445-518) |
| ***IL2RG*** | 1-2 (1-45), 2-8 (51-340), 8 (352-370) |
| ***IL7R*** | 5-7 (180-292) |
| ***JAK1*** | 3-22 (3-1023), 22-24 (1026-1123) |
| ***JAK2*** | 10 (405-442), 12-14 (505-622), 16 (665-711), 18 (762-812) |
| ***JAK3*** | 2-23 (1-1069) |
| ***KDM6A*** | 1-29 (1-1402) |
| ***KIT*** | 8-9 (411-514), 11 (550-592), 17 (788-828) |
| ***KMT2A*** | 2 (145-168), 3-4 (176-1075), 4 (1081-1112), 5 (1117-1184), 6 (1190-1212), 7 (1224-1325),  8-13 (1338-1560), 14-15 (1566-1665),  27 (2186-2195), 27 (2201-2355), 27 (2373-3215), 27 (3223-3324), 27 (3339-3575) |
| ***KRAS*** | 2-4 (1-150) |
| ***MAP2K1*** | 2 (27-90), 3 (98-146) |
| ***MPL*** | 10 (490-522), 12 (552-636) |
| ***NF1*** | 2-5 (21-189), 6 (201-218), 8-13 (244-467), 13-24 (478-1066), 25-26 (1082-1146),  26-31 (1160-1378), 31-35 (1380-1550), 35-38 (1564-1868),  39 (1870-1884), 39-47 (1886-2322), 47-52 (2325-2555), 52-58 (2568-2840) |
| ***NOTCH1*** | 26-28 (1529-1795), 34 (2069-2273), 34 (2290-2556), 34 (2069-2273), 34 (2290-2556) |
| ***NPM1*** | 11 (283-295) |
| ***NRAS*** | 2-4 (1-150) |
| ***PAX5*** | 1-10 (14-392) |
| ***PHF6*** | 2-10 (1-366) |
| ***PIGA*** | 2 (1-6), 2-6 (16-485) |
| ***PML*** | 3 (201-255) |
| ***PRPF40B*** | 2-19 (2-609), 19-20 (611-658), 20-26 (661-893) |
| ***PTEN*** | 7-8 (212-285), 8 (290-342) |
| ***PTPN11*** | 3-4 (46-125), 7 (253-285), 12 (460-462), 12-13 (465-533) |
| ***RAD21*** | 2-14 (1-632) |
| ***RARA*** | 6-7 (211-338) |
| ***RUNX1*** | 2-9 (1-437), 9 (456-474) |
| ***SETBP1*** | 4 (838-885) |
| ***SF1*** | 1-13 (1-640) |
| ***SF3A1*** | 1-7 (1-322), 7-9 (328-424), 9-16 (427-794) |
| ***SF3B1*** | 13-16 (574-790) |
| ***SH2B3*** | 2 (1-118), 2 (132-164), 2-8 (211-576) |
| ***SMC1A*** | 1-19 (1-983), 20-25 (992-1234) |
| ***SMC3*** | 1-6 (1-110), 6-16 (113-504), 16-17 (507-580), 17-29 (591-1217) |
| ***SRSF2*** | 1 (1-38), 1 (45-121) |
| ***STAG1*** | 2 (1-5), 3-20 (10-703), 21-22 (718-738), 22-27 (740-953), 27-34 (955-1259) |
| ***STAG2*** | 2-15 (1-512), 16-33 (521-1232) |
| ***STAT3*** | 17 (521-534), 17 (489-503), 17 (506-508), 18-22 (534-715) |
| ***STAT5A*** | 3-7 (1-214), 8-9 (264-286), 9-20 (303-795) |
| ***STAT5B*** | 16 (636-693) |
| ***SUZ12*** | 1-2 (20-107), 4-5 (129-169), 7-16 (198-740) |
| ***TERT*** | 1 (1-24), 2 (80-172), 2-4 (258-630), 4-5 (633-677), 5-6 (683-749), 6-8 (753-800), 8-16 (805-1133) |
| ***TET2*** | 3 (1-77), 3 (91-826), 3 (829-853), 3-11 (867-2003) |
| ***TP53*** | 2 (1-25), 4-11 (80-394) |
| ***U2AF1*** | 2 (15-44), 6 (117-161) |
| ***U2AF2*** | 1-5 (1-161), 6-12 (163-473) |
| ***WT1*** | 1 (122-216), 1 (2-44), 1 (56-58), 2-10 (216-518) |
| ***ZRSR2*** | 1-4 (1-90), 5 (108-131), 6-9 (134-263), 9-11 (267-483) |

**Table S1. Characteristics at transformation to acute myeloid leukemia.**

| **Characteristic** | **AML (N=18) N(%)/Median [range]** |
| --- | --- |
| Age (years) | 68 [46-89] |
| Male | 45 (69) |
| WBC (x10^9^/L) | 19.7 [4.3-89.2] |
| Neutrophils (%) | 33 [8-88] |
| Promyelocytes (%) | 0 [0-0] |
| Myelocytes (%) | 0 [0-0] |
| Metamyelocytes (%) | 4 [1-33] |
| Monocytes (%) | 11 [0-34] |
| Lymphocytes (%) | 18 [3-42] |
| Basophil (%) | 1 [0-25] |
| Eosinophil (%) | 1 [0-12] |
| Blast (%) | 14 [0-77] |
| Hgb (g/dL) | 9.7 [7.3-12.7] |
| Platelets (x10^9^/L) | 43 [3-361] |
| BM blast (%) | 32 [20-73] |
| BM progranulocytes (%) | 2 [0-8] |
| BM Myelocytes (%) | 8 [0-16] |
| BM Metamyelocytes (%) | 7 [1-14] |
| BM granulocytes (%) | 15 [6-45] |
| BM eosinophils (%) | 2 [0-8] |
| BM basophils (%) | 1 [0-10] |
| BM monocytes (%) | 5 [1-25] |
| Cytogenetics* |  |
| Normal | 3 (17) |
| Trisomy 8 | 3 (17) |
| i(17q) | 4 (22) |
| Del(20q) | 1 () |
| Complex | 3 (17) |
| Monosomy 7 | 1 () |
| Other | 2 () |
| NA | 4 (22) |

*: Some patients had multiple of the noted cytogenetic abnormalities and are counted in multiple of the cytogenetic

**Table S2. Univariate and multivariate analysis for survival (P-value cutoff of 0.200)**

|  | UVA | | | Backward MVA | | |
| --- | --- | --- | --- | --- | --- | --- |
|  | P | HR | 95% CI | P-value | HR | 95% CI |
| Age | 0.176 | 1.029 | 0.987-1.073 | 0.001 | 1.107 | 1.045-1.173 |
| Female | 0.800 | 0.911 | 0.444-1.872 |  |  |  |
| PS ≥2 | 0.059 | 2.053 | 0.973-4.332 |  |  |  |
| Prior malig. | 0.534 | 1.319 | 0.551-3.153 |  |  |  |
| Prior chemo/XRT | 0.666 | 1.299 | 0.396-4.259 |  |  |  |
| Prior transfusion | 0.551 | 1.233 | 0.620-2.452 |  |  |  |
| Splenomegaly | 0.761 | 1.108 | 0.574-2.139 |  |  |  |
| B symptoms | 0.483 | 1.290 | 0.633-2.630 |  |  |  |
| TLS | 0.876 | 1.087 | 0.383-3.082 |  |  |  |
| WBC | 0.282 | 1.002 | 0.999-1.005 |  |  |  |
| Neu% | 0.498 | 0.994 | 0.977-1.011 |  |  |  |
| Blasts% | 0.192 | 1.051 | 0.975-1.132 | 0.067 | 0.889 | 0.784-1.008 |
| Mono% | 0.451 | 0.962 | 0.871-1.063 |  |  |  |
| Lymph% | 0.223 | 0.975 | 0.935-1.016 |  |  |  |
| Baso% | 0.860 | 1.027 | 0.762-1.385 |  |  |  |
| Eo% | 0.076 | 0.849 | 0.708-1.018 |  |  |  |
| Hgb | 0.033 | 0.846 | 0.726-0.987 | 0.024 | 0.784 | 0.635-0.968 |
| Plt | 0.156 | 0.998 | 0.995-1.001 | 0.003 | 0.993 | 0.988-0.997 |
| BM blasts | 0.013 | 1.089 | 1.018-1.165 | <0.001 | 1.414 | 1.223-1.635 |
| BM Eo | 0.339 | 0.932 | 0.807-1.077 |  |  |  |
| BM baso | 0.935 | 1.005 | 0.884-1.143 |  |  |  |
| BM mono | 0.027 | 1.172 | 1.018-1.350 | 0.041 | 1.215 | 1.008-1.466 |
| EPO | 0.804 | 1.000 | 0.999-1.001 |  |  |  |
| LDH | 0.065 | 1.000 | 1.000-1.000 | <0.001 | 1.000 | 1.000-1.000 |
| UA | 0.435 | 1.042 | 0.939-1.157 |  |  |  |
| Diploid | 0.130 | 0.607 | 0.318-1.158 |  |  |  |
| SCT (time dep) | 0.198 | 0.454 | 0.136-1.511 | 0.007 | 0.144 | 0.035-0.593 |
